# Combinatorial approaches using an AI/ML-driven drug development platform targeting insulin resistance, lipid dysregulation, and inflammation for the amelioration of metabolic syndrome in patients

**DOI:** 10.1101/2021.09.01.458488

**Authors:** Maksim Khotimchenko, Nicholas E. Brunk, Mark S. Hixon, Daniel M. Walden, Hypatia Hou, Kaushik Chakravarty, Jyotika Varshney

## Abstract

Dysregulations of key signaling pathways leading to metabolic syndrome (MetS) are complex eventually leading to cardiovascular events and type 2 diabetes. Dyslipidemia induces progression of insulin resistance and provokes release of proinflammatory cytokines resulting in chronic inflammation, acceleration of lipid peroxidation with further development of atherosclerotic alterations and diabetes. We have proposed a novel combinatorial approach using FDA approved compounds targeting IL-17a and DPP4 to ameliorate a significant portion of the clustered clinical risks in patients with MetS. As MetS is considered a multifactorial disorder, the treatment measures cannot be focused on the specific pathway because other metabolic changes keep the pathological processes in progression. In our present research we have modeled an outcomes of metabolic syndrome treatment using two distinct drug classes. Targets were chosen based on the clustered clinical risks in MetS; dyslipidemia, insulin resistance, impaired glucose control, and chronic inflammation. The AI/ML platform, BIOiSIM, was used in narrowing down two different drug classes with distinct mode of action and modalities. Preliminary studies demonstrated that the most promising drugs belong to DPP-4 inhibitors and IL-17A inhibitors. Alogliptin was chosen to be a candidate for regulating glucose control with long term collateral benefit of weight loss and improved lipid profiles. Secukinumab, IL-17A sequestering agent used in treating psoriasis, was selected as a candidate to address inflammatory disorders. Our analysis suggests this novel combinatorial approach has a high likelihood of ameliorating a significant portion of the clustered clinical risk in MetS.

**Author summary:** Metabolic syndrome is a global epidemic affecting a significant population worldwide. This syndrome is the manifestation of clustered clinical conditions that cannot be fully ameliorated with monotherapies. No therapeutic approaches were confirmed to be effective in deceleration of the metabolic syndrome progression. Artificial intelligence driven computation methods were used to predict efficacy of innovative combinatorial therapy using IL-17A sequestering agent and a DPP-4 inhibitor. They are expected to mitigate a significant portion of the clustered risks in metabolic syndrome disrupting key pathological pathways playing important role in development of this syndrome. The main therapeutic effects are related to reduction of the elevated lipid level and high glucose concentration. Combinatorial treatment could potentially stop or reverse a significant portion of the clinical risks in metabolic syndrome globally. Repurposing of approved FDA drugs can have increased likelihood of approval of the new therapeutic regimens and can reach patients faster with reduced costs of treatment.

## INTRODUCTION

Metabolic syndrome (MetS) is a group of clinical conditions manifested as abdominal obesity, hyperglycemia, hypertension, dyslipidemia, and chronic inflammation, concurrently leading to marked increase in the risk of heart disease, diabetes and stroke, or all three [1, 2]. Currently, MetS is a growing health concern due to its increasing prevalence globally [3]. Statistics on MetS frequency of occurrence vary depending on the subgroup (social status, ethnicity, etc.) and diagnostic criteria; however, it is estimated approximately 25% of the world population have MetS [4, 5] while around 35% to 40% of the US population has MetS. The rapid increase in obesity rates in the last decades, which promotes insulin resistance [6] has driven the prevalence of MetS. Numerous clinical studies confirm direct correlation between MetS progression and type 2 diabetes mellitus incidence [7] and faster development of atherosclerosis [8]. Medical care of MetS patients including therapeutic treatment is essential, as these patients are predisposed to a variety of cardiovascular, cerebral, and hepato-renal complications as well as increased overall mortality [9-12].

During MetS, the development of high adipose accumulation represents a risk factor for cardiovascular disease and insulin resistance [13]. Brown and white adipose tissue regulate numerous metabolic pathways that, when altered, can lead to an impaired carbohydrate and lipid metabolism. Hypertrophy and hyperplasia of the adipocytes result in metabolic alterations and in the onset of a low grade chronic inflammatory state [14]. Pro-inflammatory substances, when released from adipose tissue, can trigger insulin resistance and diabetes mellitus as well as cardiovascular disorders [15]. In MetS patients, studies have identified the presence of inflammatory cytokines, such as IL-1, IL-17, and IL-18, as playing a key role in the development of atherosclerotic alterations in blood vessels caused by lipid imbalance [16, 17]. Dyslipidemia in MetS is characterized by increased circulating triglyceride levels, reduced HDL-C concentrations and therefore, an increase in low density lipoprotein cholesterol. These alterations, in turn, are closely related to impaired glucose metabolism and chronic inflammation, leading to a feedback loop encompassing MetS [18]. The abovementioned systemic inflammatory markers are the main risk factors for the development of macrovascular complications that lead to significant increase in morbidity and mortality. Manifestation of clustered clinical conditions in MetS is one of the leading causes of liver steatosis, which is confirmed by recurrence of non-alcoholic fatty liver disease in patients with MetS after liver transplantation [19].

MetS as a whole cannot be treated by a single therapeutic. Therefore, therapeutic measures generally focus on specific sub-syndromes that present themselves in a symptomatic fashion, in particular hypertension and lipid imbalance. Generally, initial treatment focuses on the lifestyle modifications e.g. diet and exercise to drive weight loss, which is still the first-line recommendation for prevention and even treatment of MetS [20]. Lifestyle intervention resulting in a 7% weight loss drove resolution of MetS in 15.6% in participants who were followed for a mean of 3.2 years. Unfortunately, only 50% of patients can achieve a weight loss of 7% [21, 22]. However, long-term compliance with a balanced lifestyle and diet restrictions is difficult to achieve let alone maintain in the target population. Therefore, pharmaceutical interventions are directed at achieving goals of low-density lipoprotein cholesterol level, blood pressure, blood glucose level, and hemoglobin A1c. Treatment may also involve drug therapy with antihypertensives, insulin sensitizers, and/or cholesterol-lowering agents [23]. These therapeutic interventions have palliative effects that do not impact the key disorders in metabolic pathways leading to MetS progression. Thus, there exists a major unmet medical need for novel therapeutic treatments for MetS, ultimately preventing the development of cardiometabolic related morbidity and mortality.

Currently there are two main approaches under investigation for alleviating MetS progression [24]. One strategy is the ‘polypill’, a variably assembled single capsule containing a combination of drugs treating several risk factors. Although the polypill cannot be titrated for better risk factor control when used alone, its advantages include simplicity and cost reduction if generic drugs are used. A second pharmacological strategy to treat patients with several risk factors while reducing the problems associated with polypharmacy is to either develop single drugs that have multiple targets or to modulate targets that affect several risk factors [25]. Therefore, despite therapeutic mitigation of the main symptoms, such as high blood pressure and elevated LDL blood level, MetS continues to progress in the patient. Relief of symptoms without direct intervention against the clustered risks in MetS treatment leads to inevitable complications. To that end, several approaches can be used successfully, such as repurposing of FDA approved drugs for combination therapies to mitigate and potentially reverse the majority of the clustered risks in MetS in patients.

Currently, the repurposing of drugs for combinatorial therapies targeting key pathways involved in MetS progression is a path of least resistance in establishing a pipeline for effective therapeutics. Commonalities affecting redundant signaling mechanisms between the sequential polytherapy and aggregate polypill therapeutic approaches exist, promoting a strategy targeting these common pathways known to upregulate insulin resistance, lipid metabolism, and expression of pro-inflammatory cytokines. Focus on these multi-stage-related biological targets inherent to both primary pathological hypotheses of MetS development is a promising avenue for the rapid repurposing of existing therapeutics.

The data-rich nature of well-studied, FDA-approved drugs is particularly amenable to our *in silico* AI/ML-integrated modeling platform BIOiSIM. The platform enables high-throughput computational compound screening based on experimentally-validated simulations of *in vivo* pharmacokinetic-pharmacodynamic (PK-PD) phenomena. Our research strategy entails the development of novel models for PK-PD predictions from repurposed drugs leading to combination therapies, generated by an integrated AI/ML-driven in silico platform, BIOiSIM. This novel combinatorial approach targets biochemical pathways responsible for the development of insulin resistance with consequent dyslipidemia and chronic inflammation, accounting for ∼70% of the clustered clinical conditions manifesting in MetS. The key pathways include but are not limited to, signaling pathways such as farnesoid X receptors (FXR), peroxisome proliferator-activated receptor (PPAR)-α, δ, and γ, fibroblast growth factor 21 (FGF21), dipeptidyl peptidase 4 (DPP-4) regulated pathways, and IL-17a regulated inflammatory pathways. FXR and PPAR-targeted drugs were found to cause toxicity issues after long-term administration and have insufficient PK profile. DDP-4, an enzyme that degrades a group of gastrointestinal hormones, incretins, affects both mechanisms of the MetS development i.e. insulin resistance and lipid metabolism. Inhibition of this pathway leads to increased insulin sensitivity and reduction of the blood lipid level in both preclinical and clinical settings. DPP-4 inhibitors have long term collateral benefits compared to drug targeting the FXR and PPAR signaling axis [26, 27]. Hence, we selected the DPP-4 inhibitor Alogliptin as a candidate for our investigation. Among the regulators of proinflammatory processes, one of the most promising cytokines is IL-17A, produced by specific CD4+ T helper (Th) cells. Dysregulation of IL-17A leads to the numerous immune-mediated and metabolic disorders. IL-17A plays a critical role in neutrophil recruitment, angiogenesis, inflammation, and autoimmune disease [28] including pulmonary, cardiac, and liver fibrosis via IL-17RA and MAPK signaling [29, 30]. Recently, IL-17A upregulation in response to diet induced obesity induces PPARγ phosphorylation adipocytes in a CDK5-dependent manner, thereby modulating diabetogenic and obesity gene expression, this correlates with IL-17A signaling in the fat of individuals with morbid obesity. Here upregulation of IL-17A and its unexpected role in adipocyte biology, in which its direct-action correlates in pathogenic reprogramming of adipocytes, causes diet induced obesity and metabolic syndrome. Hence, targeting the IL-17A signaling axis could be an effective treatment for the clustered clinical risks manifested in MetS [31]. In the current clinical landscape, a single therapy has been at best palliative, unfortunately not showing any promise in reversing any of the clustered clinical conditions of MetS. We present here a novel approach that entails a repurposing and combinatorial therapeutics approach against major clinical risks in MetS. Our approach is based on PK-PD predictions of two distinct modalities, generated by our proprietary AI/ML in silico platform BIOiSIM. By targeting dyslipidemia and inflammation in the patient population, we have the potential in ameliorating a large percentage of the clustered risks manifested in MetS and address the medical unmet needs.

In the present study, we have used AI/ML based computational modeling and simulations to generate PD predictions for novel combinatorial therapies for repurposed drugs Alogliptin and Secukinumab. These drugs target critical pathways in insulin resistance and inflammation in the progressive development of MetS. Using abundant PK-PD data from publicly available sources (e.g. clinical and preclinical datasets), we have modeled the disposition and potential dosing regimen of various FDA-approved pharmaceutical compounds in target tissues known to be involved in key stages of the MetS progression. An understanding of compounds capable of optimal target-tissue deposition and regulation of the biological target structures will help accelerate the development of a targeted anti-MetS combinatorial therapies. Overall, our investigation has identified FDA-approved therapeutics using our AI/ML driven drug development platform, for repurposing of drugs, leading to combination therapies and associated dosing regimens in the patient populations.

## RESULTS AND DISCUSSIONS

IL-17A plays an important role in the commonality that is manifested in the pathophysiology of clinical conditions in MetS. These similarities include an inflammatory presence associated with the development of modified autoantigens targeted by both arms of the immune system; the innate and adaptive immune system [37]. Th1 in both scenarios plays an initiatory catalyst. The alteration of the vesicular blood flow started by invasion of these inflammatory cells into the vessel wall is initially activated by adhesion molecules whose production is controlled by the secretion of pro-inflammatory cytokines and chemokines [37]. Historic evidence establishing the correlation between cardiometabolic disease with psoriasis has existed for a long time [38]. Severe psoriasis is associated with an increased risk of cardiovascular mortality [39], with a subsequent reduction in life expectancy in patients [40]. Recently, cumulative research evidence demonstrated that IL-17A may represent one of the main links between cardiometabolic disease manifestations and psoriatic inflammation [41, 42]. Additionally, it has been established that psoriasis patients have an increased likelihood of high body mass index [43], and have higher chances of having metabolic syndrome [44]and type 2 diabetes [41]. In the study of Bruin et al. [33], three weekly subcutaneous doses of 150 mg Secukinumab followed by subcutaneous dosing every 4 weeks had contributed to the rapid onset of psoriatic lesion clearance confirmed by decreased hBD-2 in both serum and skin lesions. Since IL-17A mediated inflammation is implicated in a segment of the clustered risk, specifically obesity, atherosclerosis and low grade chronic inflammatory disease, in metabolic syndrome we conducted a meta-analysis of the phase 3 JUNCTURE study to determine the reduction in free circulating levels of IL-17A. As seen in figure 1 left panel, our modeling of free IL-17A levels suggests that during the loading phase of Secukinumab, circulating levels of free IL-17A are reduced by 98% and during steady state dosing of 150 mg Secukinumab every 4 weeks, circulating levels of IL-17A remain reduced to less than 5% of untreated. Based on our simulated outcomes Secukinumab is a potent IL-17A inhibitor and can be repurposed in targeting a significant portion of the inflammation-driven clustered risks of MetS.

**Figure 1.**
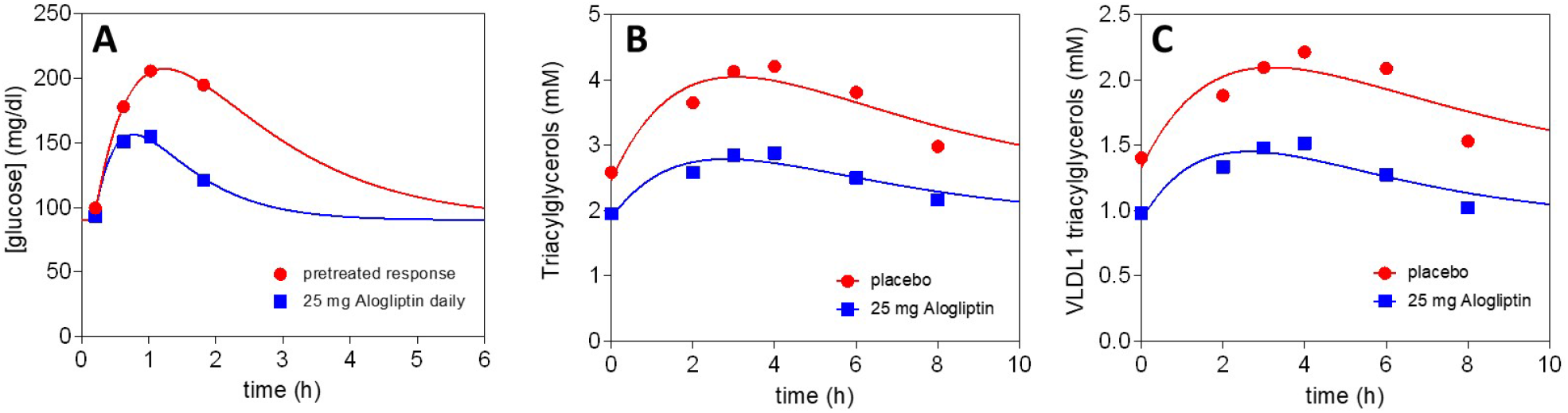
Simulations of IL-17A inhibitor Secukinumab pharmacokinetics and its engagement with IL-17A target. Secukinumab is dosed 150 mg s.c. weekly for 4 weeks then once every 4 weeks. Simulations based on published reports.

Type 2 diabetes (T2D) mellitus manifests itself as impaired blood glucose control underlying multiple metabolic abnormalities, including decreased level of insulin secretion and prolonged exposure to free fatty acids [45], insulin resistance, progressive loss of beta-cell function, impaired glucagon secretion, and dysregulation of incretin hormone signaling [46]. Alogliptin blocks dipeptidyl peptidase 4 (DPP-4) enzyme, prolonging the exposure and levels of the incretin hormones, glucagon-like peptide-1 (GLP-1) and glucose-dependent insulinotropic polypeptide (GIP) [47]. These incretin hormones increase insulin secretion and decrease glucagon secretion, which also decreases hepatic glucose production and reduction in free fatty acid and or lipid levels [47]. Alogliptin is commonly prescribed at 25 mg daily for T2D, and our meta-analysis of the report by Christopher et al. [26] reveals DPP-4 is inhibited by 90.5% (daily through = 84%, daily peak = 96.8%) at steady state. Under this treatment regimen, projections of the simulated dose would lead to fasting glucose levels reduction by an average of 9% and triacylglycerides by an average of 10%. Long term administration of alogliptin has shown global reductions in glucose levels and improved plasma lipid profile [46]. In addition, the agent is relatively well tolerated with few adverse effects [45]. A meta-analysis of the glucose challenge test presented in figure 1 of [36] suggests a daily dose of 25 mg Alogliptin increases patient systemic glucose clearance by approximately 3-fold.

In addition to enhanced blood glucose regulation, in a report by Aramaki et al. [36] Aloglipitin reduces postprandial triacylglycerol levels. A meta-analysis of the figure 1 data provided by aforementioned article indicates daily doses of 25 mg Aloglipitin produces modest basal reductions in blood triacylglycerol levels and reduces the overall triacylglycerol exposure. Basic pharmacodynamic modeling indicates the rates of lipid absorption and clearance from blood are unaffected; rather, it seems the absorption from the gut is reduced or the liver captures more in the first pass.

MetS covers a group of clustered clinical conditions leading to T2D and cardiovascular events in patients. These clinical conditions, if left unchecked, present several risks that could lead to high morbidity and mortality. Therefore, accelerating the development of therapeutic approaches is critical to patients. To date, relatively unsuccessful monotherapies or combinations therapies specifically using small molecules as a modality have been used in patients in hope of stopping and/or reversing a single clinical condition in the gamut of conditions encompassing MetS. We have investigated a novel approach using our AI/ML-driven BIOiSIM platform to accelerate the drug development process for a combination therapy using repurposed compounds consisting of two different modalities and mechanisms of action. Among the many FDA approved drug classes targeting different clinical conditions spanning the spectrum ranging from insulin resistance, dyslipidemia and inflammation under MetS, we have chosen a small molecule, Alogliptin (DPP-4 inhibitor) and Secukinumab (anti-IL-17a antibody) for a potential combinatorial approach targeting and possibly ameliorating a significant portion of MetS. This approach takes advantage of two distinct mechanisms of action cumulatively leading to immediate improvement resulting in initial decrease in insulin resistance and improved glucose control. This results in long-term beneficial reduction in inflammation with consequent drop in lipid levels, which may in turn result in body weight reduction. Our simulation demonstrates high potency for Alogliptin and Secukinumab in downregulating their targets as shown in Figures 2 and 3.

**Figure 2.**
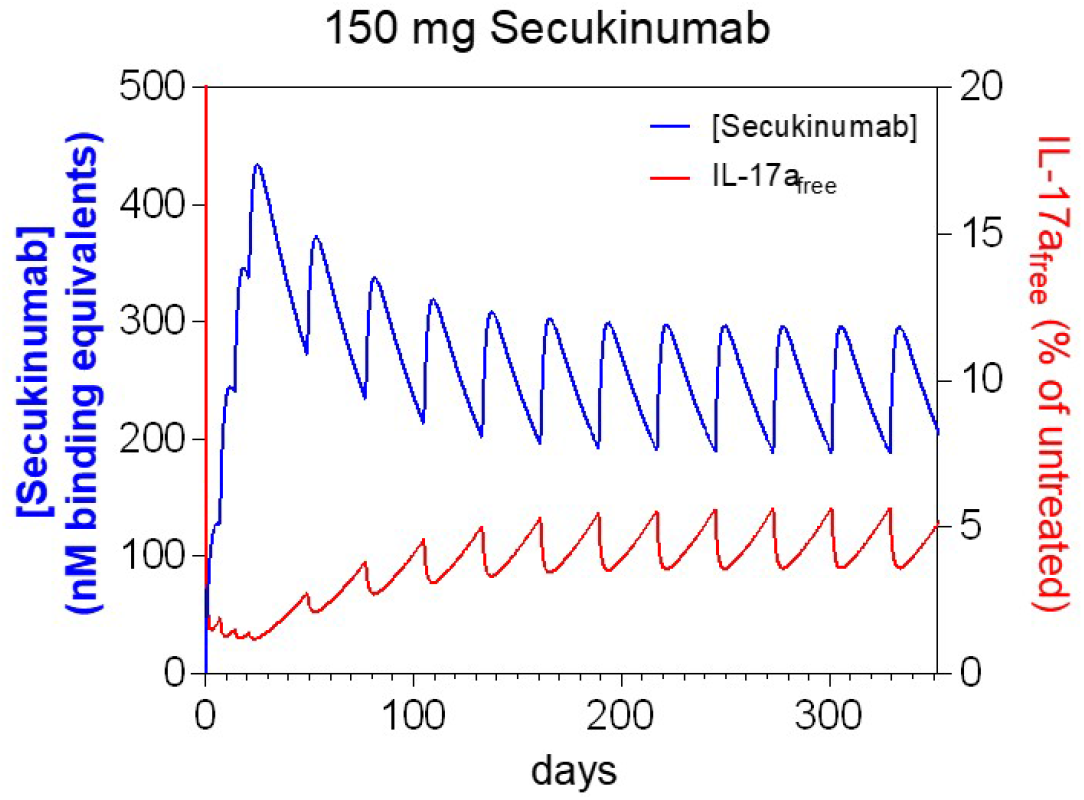
Simulations of DPP-4 inhibitor Alogliptin pharmacokinetics and its engagement with DPP-4 protein following daily 25 mg oral administration. Simulations based on published reports.

**Figure 3.**
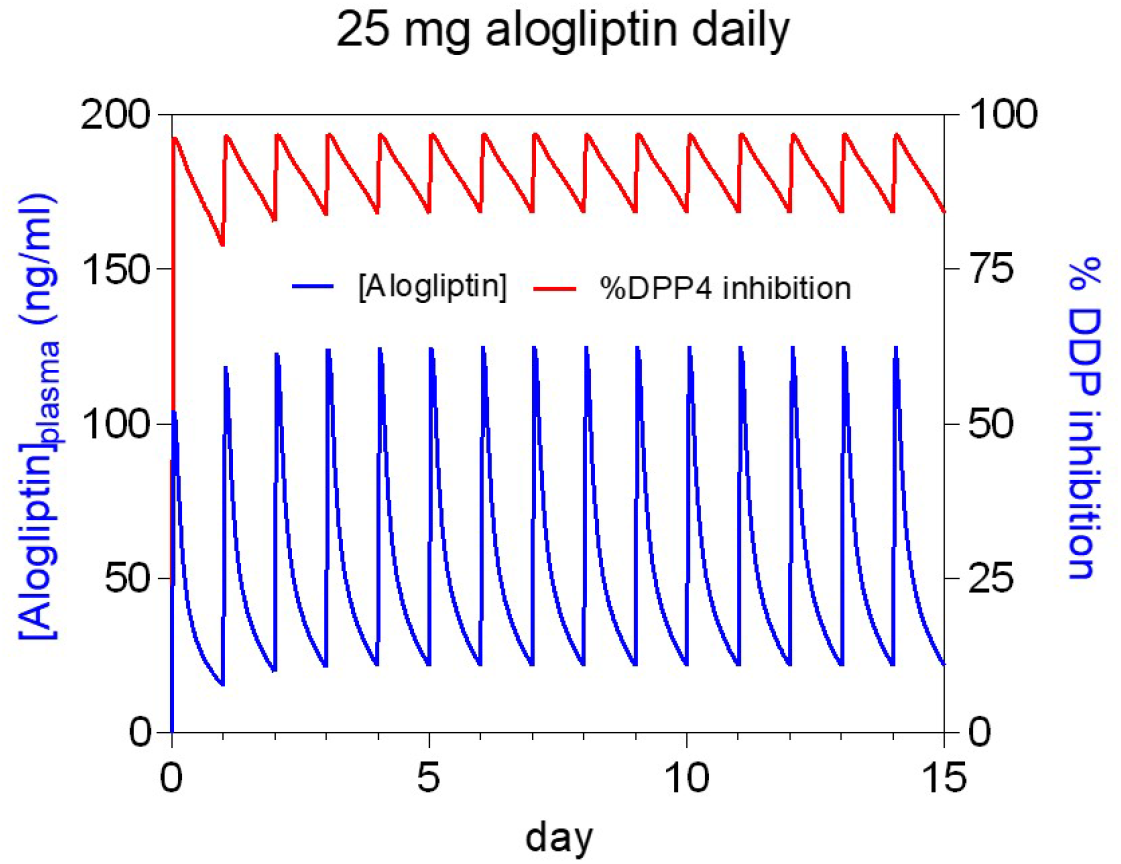
Retrospective analysis of Alogliptin impact on glucose and triacylglyceride levels. (A) Blood glucose response following 75 g glucose pre-treatment with daily 25 mg Alogliptin vs placebo (Aramaki et al., 2017). Lipid responses to 25 mg Alogliptin daily vs placebo for postprandial (B) triacylglycerol and (C) VLDL1 levels.

We therefore anticipate that prolonged exposure to the combinations of DPP-4 inhibitor and anti-IL71a therapy could reduce markedly cardiovascular events and T2D in patients with MetS.

## CONCLUSIONS

Key metabolic pathway dysregulations leading to MetS in part originate from impaired glucose control and disorders in lipid metabolism. This leads to elevated blood lipid level resulting in insulin resistance and chronic inflammation due to the release of proinflammatory cytokines. As MetS is a multifactorial-multidimensional disorder, the treatment measures cannot solely focus on the specific pathway because other metabolic changes would not be remedied, allowing progression of other problematic pathological states. Our results suggest that simultaneously targeting lipid metabolic pathways, impaired glucose control, insulin resistance, and chronic inflammation with DPP-4 inhibitor Alogliptin and IL-17A inhibitor Secukinumab will likely provide a high likelihood of ameliorating a significant portion of the clustered clinical risk associated with MetS.

## METHODS

### Overview of the BIOiSIM platform

The core functionality of our *in silico* simulation platform used in the present study was described previously [32]. Briefly, the platform is capable of utilizing AI-ML algorithms to either complete the missing parameters if the drug properties are not provided, or as predictive solutions to train existing in vivo and/or in vitro datasets. The platform performs both PBPK predictions using a 16-compartment model and PD and efficacy simulations via auxiliary models integrated into the centralized framework. The software systems are hosted on Amazon Web Services (AWS) cloud enabling high throughput, parallelized PK or PD and Efficacy simulations, which are available on the VeriSIM Life customer portal, BIOiWARE. The drug-dependent parameters used in the model are either determined in experimental settings or predicted/optimized using a combination of ML cost minimization algorithms via integrating iterative optimization algorithms with random walk methods (similar to Markov Chain/Monte Carlo) to converge on a global minimum.

### Meta-analysis of published reports

Briefly, in vivo plasma concentration datasets with related biomarker levels were manually digitized from source publications using “WebPlotDigitizer” version 4.2.34. Model development and validation was done using our in-house platform, which is primarily written in Python with Cython integration; auxiliary packages matplotlib (v2.0.2) and Numpy (v1.14.2) are also integrated into the system for simulation deployment and analysis.

### Drug PD and efficacy modeling

Secukinumab pharmacokinetic and pharmacodynamic data (serum total IL-17A levels versus time) taken from the study of Bruin et al. [33] were digitized and analyzed. Free serum concentrations of IL-17A were computed implicitly via the Morrison equation from a solution to Equation 1 [34]. All concentrations were determined in nM and Secukinumab was assumed to bind two equivalents of IL-17A. Untreated free levels of IL-17A were below the limit of quantification in the report. A 52-week study in which Secukinumab was dosed subcutaneously (s.c.) at 150 mg weekly for the first 4 weeks and then once every 4 weeks for up to 52 weeks (extracted from Bruin et al., figure 5) was used, from which total IL-17A data were used to find best fit values of the untreated IL-17A concentration and its zero-order formation rate.

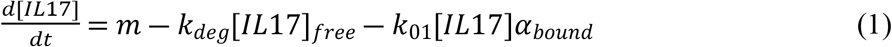

where:

*m* is the zero-order formation rate of IL-17A

[*IL*17]_*free*_ is Secukinumab bound IL-17A

*k*_*01*_ = the elimination rate constant of Secukinumab

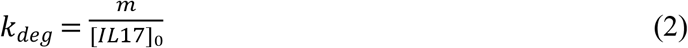

[*IL*17]_0_ is the untreated serum concentration of IL-17A,, which was assumed to have been at steady-state prior to treatment.

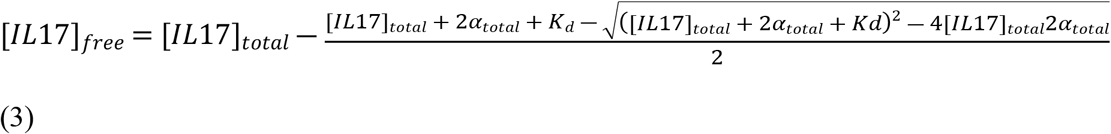

*where K*_*d*_ = 45 nM [33].

Alogliptin pharmacokinetic profile, DPP-4 activity levels and blood glucose levels after chronic dosing were extracted from [35]. Triacylglycerol levels versus Alogliptin treatment were extracted from [36]. The percent inhibition of DPP-4 was computed from a best fit of Equation 4.

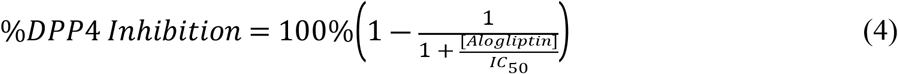

Glucose challenge test analysis, study data reported in [36] were digitized and Glucose levels were modeled assuming a basal glucose level, a rate of systemic entry (*k*_01_) and a rate of elimination (*k*_10_) and an amplitude factor (*A*) by use of equation 5. From the study data, basal glucose levels were unchanged between daily 25 mg Alogliptin treatment and pretreatment levels. Pre-treatment and treated glucose profiles were fit globally with all parameters common to cohorts except glucose clearance (*k*_10_).

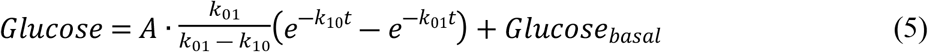

Triglyceride level analysis, study data reported in [27] were digitized and lipid levels were modeled initially by use of Equation 5 but it became apparent from fitting that the rate into plasma was equal to the rate of elimination (*k*_01_ = *k*_10_), which reduces the system to Equation 6. DPP-4 inhibition had an influence on all three descriptive parameters.

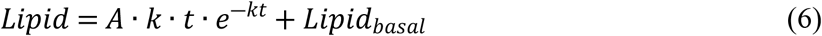

## AUTHOR CONTRIBUTIONS

**Conceptualization:** Maksim Khotimchenko, Kaushik Chakravarty

**Data curation:** Maksim Khotimchenko, Hypatia Hou

**Formal analysis:** Maksim Khotimchenko

**Investigation:** Mark S. Hixon, Maksim Khotimchenko

**Methodology:** Mark S. Hixon

**Supervision:** Jyotika Varshney

**Validation:** Daniel M. Walden

**Visualization:** Nicholas E. Brunk, Mark S. Hixon

**Writing – original draft:** Mark S. Hixon, Maksim Khotimchenko

**Writing – review & editing:** Kaushik Chakravarty, Jyotika Varshney

